# SON-mediated circular RNA suppression promotes PKR signaling and radiation resistance in glioblastoma

**DOI:** 10.64898/2026.09.28.755153

**Authors:** Caitlin A. Harvey, Sophia Dunlap, Mostafa Mohamed, Eshika Kudaravalli, Amr Elkholy, Leah Gehrs, Saeed Zakakhosravi, Bohye Park, Kwang Wook Min, James M. Murphy, Opeyemi Iwaloye, Satoru Osuka, Ssang-Taek Steve Lim, Christopher D. Willey, Erik K. Flemington, Eun-Young Erin Ahn

## Abstract

Glioblastoma (GBM) exhibits extensive loss of circular RNAs (circRNAs), yet the mechanisms driving this depletion and its functional consequences remain unclear. Here, we identify the splicing co-factor SON as a previously unrecognized suppressor of circRNA biogenesis in GBM. CircRNA sequencing revealed widespread increases in circRNA abundance following SON knockdown. Conversely, elevated SON expression in GBM, with further increases in radiation-resistant models, was associated with widespread circRNA loss, including reduced expression of circUSP1 and circSUCO. SON knockdown increased circRNA abundance and attenuated activation of the dsRNA-responsive kinase PKR and downstream NFκB signaling, whereas radiation-resistant GBM cells exhibited enhanced PKR/NFκB activation that was attenuated by SON depletion. Importantly, overexpression of the SON-suppressed circRNAs circUSP1 and circSUCO reduced PKR activation and selectively decreased clonogenic survival following irradiation in radiation-resistant GBM cells, with little effect on parental cells. Together, these findings identify a previously unrecognized SON/circRNA/PKR regulatory pathway linking RNA processing to acquired radiation resistance in GBM and highlight specific PKR-inhibitory circRNAs as potential therapeutic tools for resensitizing radiation-resistant GBM to radiotherapy.

## Introduction

Glioblastoma (GBM) is the most aggressive primary brain tumor and remains one of the deadliest human cancers (1). Despite maximal surgical resection followed by temozolomide chemotherapy and radiotherapy, the median survival time is only 14 - 15 months after diagnosis (2). Nearly all tumors recur (3, 4), reflecting the remarkable capacity of GBM cells to survive cytotoxic stress. A major driver of treatment failure and recurrence is the emergence of radiation resistance, a phenotype shaped by strong intratumoral heterogeneity and an immunosuppressive microenvironment (5). Understanding the molecular mechanisms that enable GBM cells to survive cytotoxic stress remains critical for improving patient outcomes.

Circular RNAs (circRNAs) are covalently closed RNA molecules formed by backsplicing, a process in which a downstream 5’ splice site is joined to an upstream 3’ splice site (6). CircRNAs are particularly abundant in the brain compared with other tissues and play an integral role in neuronal development (7), but they have also been implicated in various cancers (8–12). Notably, GBM exhibits lower circRNA expression than normal brain tissue (13, 14). Because circRNA abundance increases with neuronal differentiation and decreases in neural stem-like states (7), the loss of circRNAs in GBM may reflect the poorly differentiated phenotype that characterizes these tumors.

An emerging function of circRNAs is the regulation of innate immune signaling, including modulation of protein kinase R (PKR), a stress-responsive kinase activated by double-stranded RNA (dsRNA), growth factors, cytokines, and cellular stress (15–17). PKR activity is elevated in multiple cancers, including GBM (18–21). Activated PKR promotes NFκB signaling through interaction with IKK (22), and NFκB is a well-established driver of GBM progression and therapeutic resistance (23). Consistent with this role, phosphorylated PKR has been implicated in resistance to chemotherapy and radiotherapy (24, 25). Importantly, many circRNAs contain short, imperfect dsRNA regions, typically less than ∼33 bp in length, that bind PKR without activating it, thereby functioning as endogenous decoys that sequester and inhibit PKR (15, 26). These findings suggest that the loss of circRNAs in GBM, particularly in radiation-resistant tumors, may relieve inhibition of PKR and promote adaptive stress-response signaling.

The mechanisms that downregulate circRNA expression in GBM remain poorly understood. SON is a nuclear speckle-associated RNA-/DNA-binding protein that regulates alternative RNA splicing. SON promotes the splicing of transcripts containing “weak” splice sites through the recruitment of spliceosomal machinery and regulates oncogenic alternative splicing (27, 28). SON is upregulated in GBM and is required for tumor growth (27), yet whether SON regulates backsplicing and circRNA biogenesis has not been explored. Given its established role in spliceosome recruitment, we hypothesized that SON may regulate circRNA biogenesis and influence PKR activity within radiation-resistant GBM.

Here, we identify SON as a previously unrecognized suppressor of circRNA biogenesis in GBM and show that SON-dependent circRNA regulation is associated with altered PKR and NFκB signaling. We further show that overexpression of either SON-suppressed circUSP1 or circSUCO inhibits PKR activation and selectively resensitizes radiation-resistant GBM cells to irradiation. Together, these findings identify a previously unrecognized SON/circRNA/PKR regulatory pathway linking RNA processing to radiation resistance and identify specific circRNAs as potential therapeutic tools for restoring radiation sensitivity in resistant GBM.

## Results

### SON knockdown increases circRNA abundance in glioma stem cells

To examine whether SON regulates circRNA expression in GBM, we performed circRNA sequencing in GSC83 glioma stem cells (29) transduced with a SON-targeting shRNA (shSON) or a control lentivirus (shCtrl). Total RNA was treated with RNase R to enrich circRNAs by degrading linear RNAs before sequencing (**Figure 1A**). Backsplice junctions were analyzed using CIRCexplorer2 (30) and CIRI2/CIRIquant (31, 32) (**Figure 1B**).

**Figure 1.**
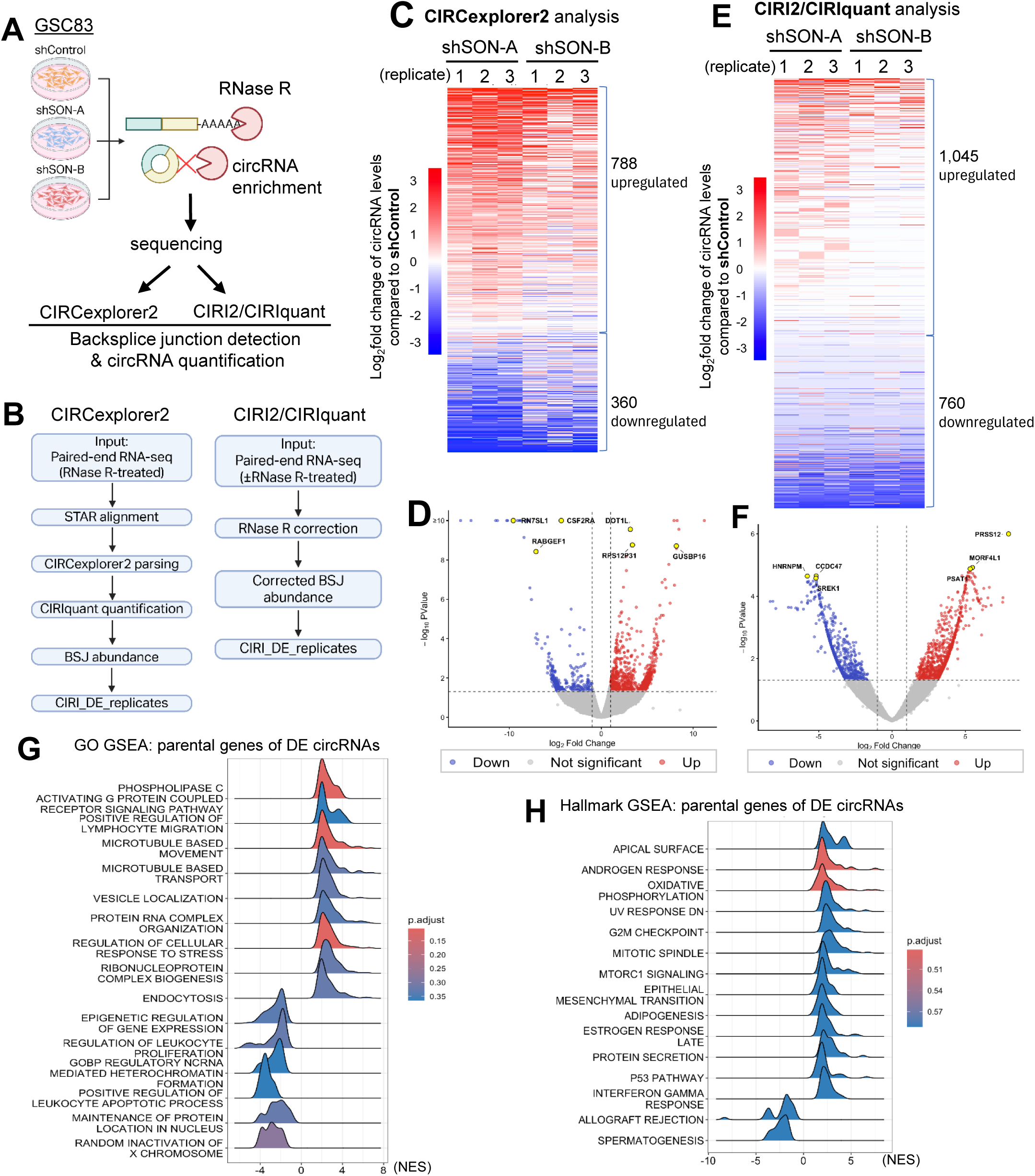
SON knockdown induces widespread circRNA upregulation in GSC83 glioma stem cells. **(A)** Schematic of the experimental workflow for circRNA sequencing in control (shCtrl) and SON knockdown (shSON) GSC83 cells. Total RNA was treated with RNase R to enrich circRNAs prior to sequencing. **(B)** Bioinformatic workflows for circRNA identification and quantification using CIRCexplorer2 and CIRI2/CIRIquant. **(C, D)** CIRCexplorer2 analysis of circRNAs in shSON relative to shCtrl cells, shown as a heatmap **(C)** and volcano plot **(D)**. A total of 788 circRNAs were increased and 360 were decreased following SON knockdown. **(E, F)** CIRI2/CIRIquant analysis of circRNAs in shSON relative to shCtrl cells, shown as a heatmap **(E)** and volcano plot **(F)**. A total of 1,045 circRNAs were increased and 760 were decreased following SON knockdown. For **(C – F)**, differentially expressed circRNAs were defined by |log_2_ fold change| > 1 and nominal *p* < 0.05. **(G)** Gene Ontology (GO) enrichment analysis of host genes associated with differentially expressed circRNAs. **(H)** Hallmark gene set enrichment analysis of host genes associated with differentially expressed circRNAs.

Both pipelines identified more increased circRNAs than decreased following SON knockdown. CIRCexplorer2 identified 788 increased and 360 decreased circRNAs, whereas CIRI2/CIRIquant identified 1,045 increased and 760 decreased circRNAs (absolute log_2_ fold change > 1; nominal P < 0.05; **Figure 1C - F**). Both analyses therefore showed a broad increase in circRNA abundance following SON depletion.

To characterize the host genes of the altered circRNAs, we performed gene set enrichment analyses. Gene Ontology terms associated with spliceosomal assembly and cellular stress response were enriched (**Figure 1G**). Hallmark analysis of host genes associated with increased circRNAs identified oxidative phosphorylation, G2/M checkpoint, and p53 signaling gene sets (**Figure 1H**). Notably, several of the identified pathways, including oxidative phosphorylation, glycolysis, and DNA damage response pathways, have previously been implicated in glioma radioresistance (33–35). These enrichment results reflect the biological pathways associated with the host genes from which the altered circRNAs are derived and do not indicate changes in pathway activity. Together, these data identify SON as a suppressor of circRNA biogenesis in GBM and show that SON depletion leads to widespread increases in backsplicing.

### Elevated SON expression in GBM is associated with suppression of SON-regulated circRNAs

To further explore the relationship between SON and circRNA biogenesis, we first confirmed the overexpression of SON in GBM compared to normal brain. Western blot analysis of normal human astrocytes (NHA), GSC83 glioma stem cells, established GBM lines (LN229 and MGG18), and patient-derived xenolines (JX14P and JX39P) revealed markedly elevated SON protein expression across all GBM models relative to NHA controls (**Figure 2A**). Together with our SON knockdown results, these findings suggest that elevated SON contributes to circRNA suppression in GBM.

**Figure 2.**
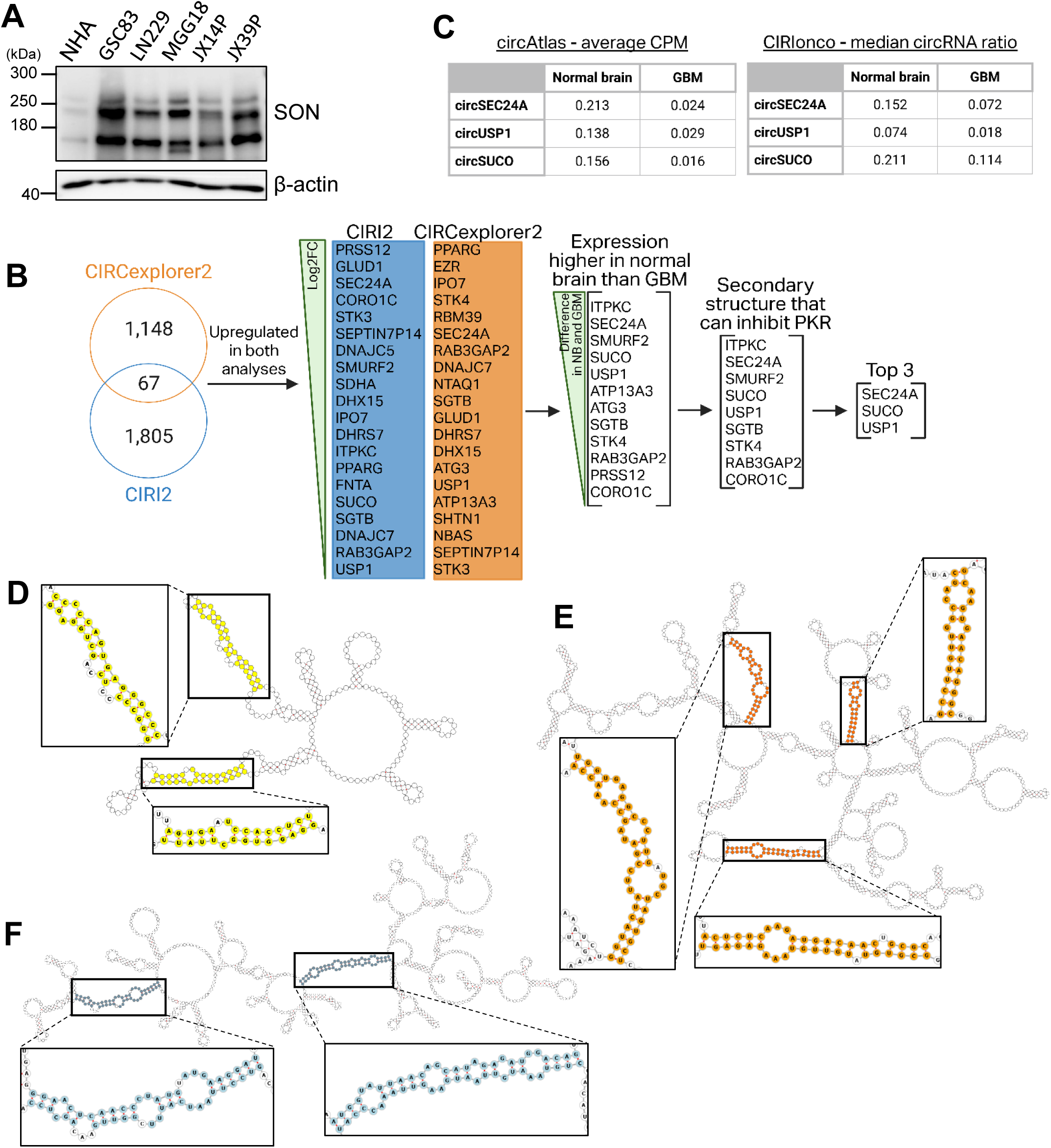
Elevated SON expression in GBM is associated with suppression of SON-regulated circRNAs. **(A)** Western blot showing elevated SON protein expression across multiple GBM models compared with normal human astrocytes (NHA). **(B)** Schematic of the multistep candidate-selection strategy used to identify SON-regulated circRNAs. CircRNAs increased following SON knockdown in both CIRI2/CIRIquant and CIRCexplorer2 analyses were ranked by log_2_ fold change and further prioritized based on differential expression between normal brain and GBM and predicted short dsRNA regions. **(C)** Expression of circSEC24A, circUSP1, and circSUCO in normal brain and GBM based on data from CIRIonco and circAtlas. **(D - F)** RNAfold-predicted secondary structures of circSEC24A **(D)**, circUSP1 **(E)**, and circSUCO **(F)**, with predicted short dsRNA regions highlighted.

We next sought to identify specific circRNAs regulated by SON in GBM (**Figure 2B**). We compared circRNAs significantly upregulated following SON knockdown in both CIRI2/CIRIquant and CIRCexplorer2 analyses and ranked the shared circRNAs by log_2_ fold change. To prioritize circRNAs with potential biological relevance to GBM, we used CiriOnco (36) and circAtlas (37) to determine which candidates showed higher expression in normal brain compared to GBM (**Figure 2C**). Given the ability of circRNAs to inhibit PKR through short dsRNA structures (15), candidate circRNAs were analyzed using RNAfold from the ViennaRNA Package (38) to identify predicted dsRNA regions of 14 - 29 bp (**Figure 2D - F**). Based on these criteria, circSEC24A, circUSP1, and circSUCO were selected for further investigation (**Figure 3A**).

**Figure 3.**
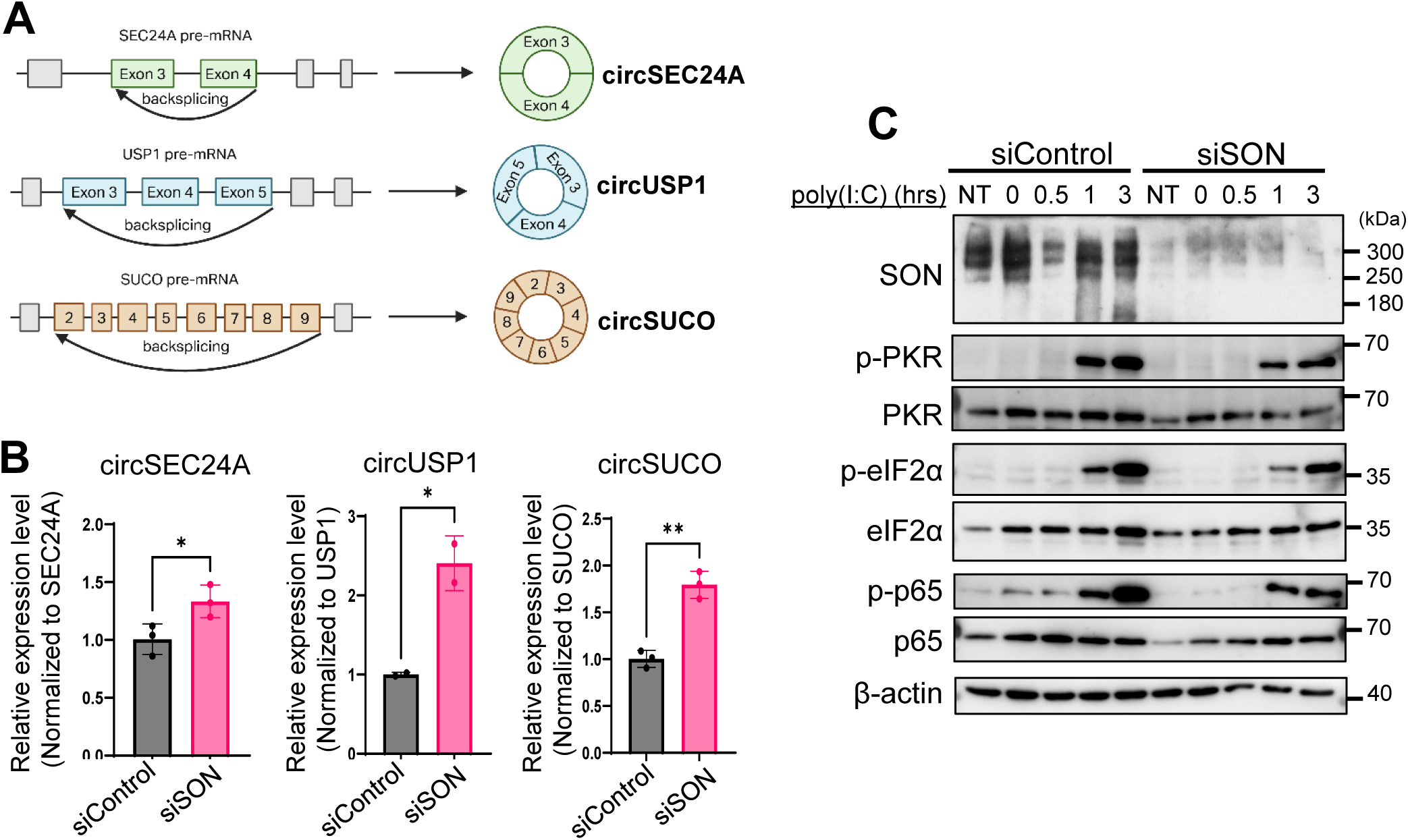
SON knockdown increases SON-regulated circRNAs and attenuates PKR activation and downstream signaling. **(A)** Schematic of the pre-mRNA exon organization and backsplicing events generating circSEC24A, circUSP1, and circSUCO. **(B)** Relative expression of circSEC24A, circUSP1, and circSUCO following SON knockdown in LN229 cells. CircRNA expression was normalized to the corresponding linear parental transcript. \**p* < 0.05; \*\**p* < 0.01; \*\*\**p* < 0.001. **(C)** LN229 cells were transfected with control siRNA (siCtrl) or SON-targeting siRNA (siSON) and stimulated with poly(I:C) for the indicated times. Western blot analysis shows SON depletion and reduced phosphorylation of PKR, p65, and eIF2α following poly(I:C) stimulation in siSON cells compared with siCtrl cells. Cells were serum-starved for 6 h before poly(I:C) stimulation, except for the non-treated (NT) group.

To validate the circRNA-sequencing results, we performed qRT-PCR using convergent primers (detecting linear and circular transcripts) and divergent primers (specific to circRNA). Upon SON knockdown using siRNA (siSON), LN229 cells exhibited increased expression of all three circRNAs when normalized to their corresponding linear transcripts (**Figure 3B**).

Together, these results identified circSEC24A, circUSP1, and circSUCO as SON-regulated circRNAs that are expressed at lower levels in GBM and contain predicted short dsRNA regions.

### SON knockdown attenuates PKR activation and downstream signaling

Having identified SON-regulated circRNAs with predicted short dsRNA regions, we next asked whether SON depletion affects PKR activation and downstream signaling. LN229 cells were transfected with siSON or control siRNA and then, after 2 days, stimulated with poly(I:C), a synthetic dsRNA used to activate PKR. Western blot analysis revealed that SON knockdown attenuated PKR activation, with reduced PKR phosphorylation at 1 and 3 hours following poly(I:C) stimulation (**Figure 3C**). Consistent with reduced PKR activation, phosphorylation of p65, indicative of NFκB activation, and phosphorylation of eIF2α were also diminished following SON depletion.

Together, these results show that SON depletion, which increases the abundance of SON-regulated circRNAs, is accompanied by attenuated PKR activation and downstream NFκB and eIF2α signaling. These findings establish an association between SON-dependent circRNA regulation and PKR signaling in GBM and provide the basis for investigating whether SON-regulated circRNAs directly mediate PKR inhibition.

### Radiation-resistant GBM exhibits elevated SON expression and global circRNA loss

Since radiation resistance remains a major barrier to the effective treatment of GBM, we next examined whether SON expression and circRNA abundance are altered during the acquisition of radiation resistance. To investigate molecular changes associated with acquired radiation resistance, we generated a radiation-resistant LN229 cell line (LN229RR) by exposing parental LN229 cells (LN229P) to 5 Gy irradiation every 4 days for four cycles (**Figure 4A**). We then compared SON expression between LN229P and LN229RR cells. Western blotting and immunofluorescence revealed increased SON expression in LN229RR compared with LN229P (**Figure 4B** and **C**). We next examined the paired patient-derived xenoline model JX39P and its radiation-resistant derivative, JX39P-RT (**Figure 4D**) (39). Consistent with the LN229 model, SON expression was increased in JX39P-RT compared with JX39P (**Figure 4E** and **F**).

**Figure 4.**
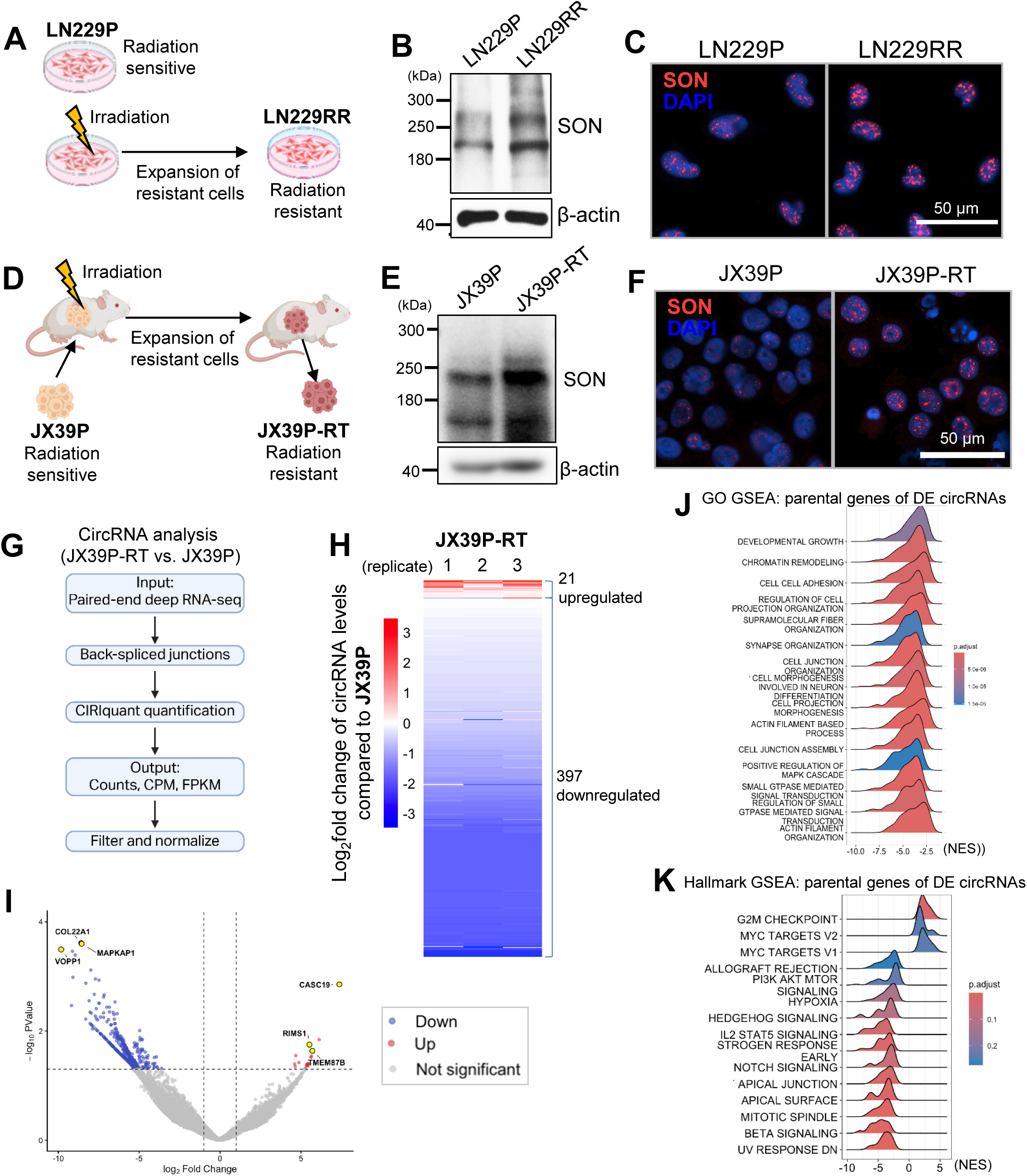
Radiation-resistant GBM exhibits elevated SON expression and widespread circRNA loss. **(A)** Schematic of the generation of the radiation-resistant LN229 model. LN229RR cells were generated from parental LN229 cells (LN229P) by repeated irradiation. **(B)** Western blot showing increased SON protein expression in LN229RR compared with LN229P cells. **(C)** Immunofluorescence staining for SON (red) in LN229P and LN229RR cells. Nuclei were counterstained with DAPI (blue). Scale bar, 50 μm. **(D)** Schematic of the paired patient-derived JX39P and radiation-resistant JX39P-RT model. **(E)** Western blot showing increased SON protein expression in JX39P-RT compared with JX39P cells. **(F)** Immunofluorescence staining for SON (red) in JX39P and JX39P-RT cells. Nuclei were counterstained with DAPI (blue). Scale bar, 50 μm. **(G)** Bioinformatic workflow for circRNA identification and quantification in JX39P and JX39P-RT using CIRI2/CIRIquant. **(H, I)** Heatmap **(H)** and volcano plot **(I)** showing widespread reduction in circRNA abundance in JX39P-RT relative to JX39P, with 397 circRNAs decreased and 21 increased. Differentially expressed circRNAs were defined by |log_2_ fold change| > 1 and nominal *p* < 0.05. **(J)** Gene Ontology enrichment analysis of host genes associated with differentially expressed circRNAs. **(K)** Hallmark gene set enrichment analysis of host genes associated with differentially expressed circRNAs.

Given our finding that SON depletion increases circRNA abundance, we next asked whether the increased SON expression observed in radiation-resistant GBM is accompanied by circRNA loss. Using RNA-sequencing data from JX39P and JX39P-RT, we performed circRNA analysis using CIRI2/CIRIquant (**Figure 4G**). JX39P-RT exhibited widespread reduction in circRNA abundance relative to JX39P, with 397 circRNAs decreased and only 21 increased (absolute log_2_ fold change > 1; nominal P < 0.05; **Figure 4H** and **I**).

To characterize the host genes associated with altered circRNAs in radiation-resistant cells, we performed gene set enrichment analysis (**Figure 4J** and **K**). Enriched gene sets included G2/M checkpoint, MYC targets, and inflammatory response pathways, with some overlap with the host-gene programs identified following SON knockdown. As in the analysis of SON-depleted cells, these enrichment results describe the host genes from which the altered circRNAs are derived and do not indicate changes in the activity of these pathways.

Together, these findings show that increased SON expression in radiation-resistant GBM is accompanied by widespread circRNA loss, consistent with the inverse relationship between SON expression and circRNA abundance observed in our earlier experiments.

### Elevated SON contributes to enhanced PKR/NFκB activation in radiation-resistant GBM

To determine whether the SON-regulated circRNAs identified following SON knockdown were also altered in radiation-resistant GBM, we measured circSEC24A, circUSP1, and circSUCO expression in the JX39P/JX39P-RT pair. qRT-PCR showed that circUSP1 and circSUCO were significantly decreased in JX39P-RT compared with JX39P, whereas circSEC24A was unchanged (**Figure 5A**). These findings indicate that a subset of SON-regulated circRNAs is further suppressed in radiation-resistant GBM.

**Figure 5.**
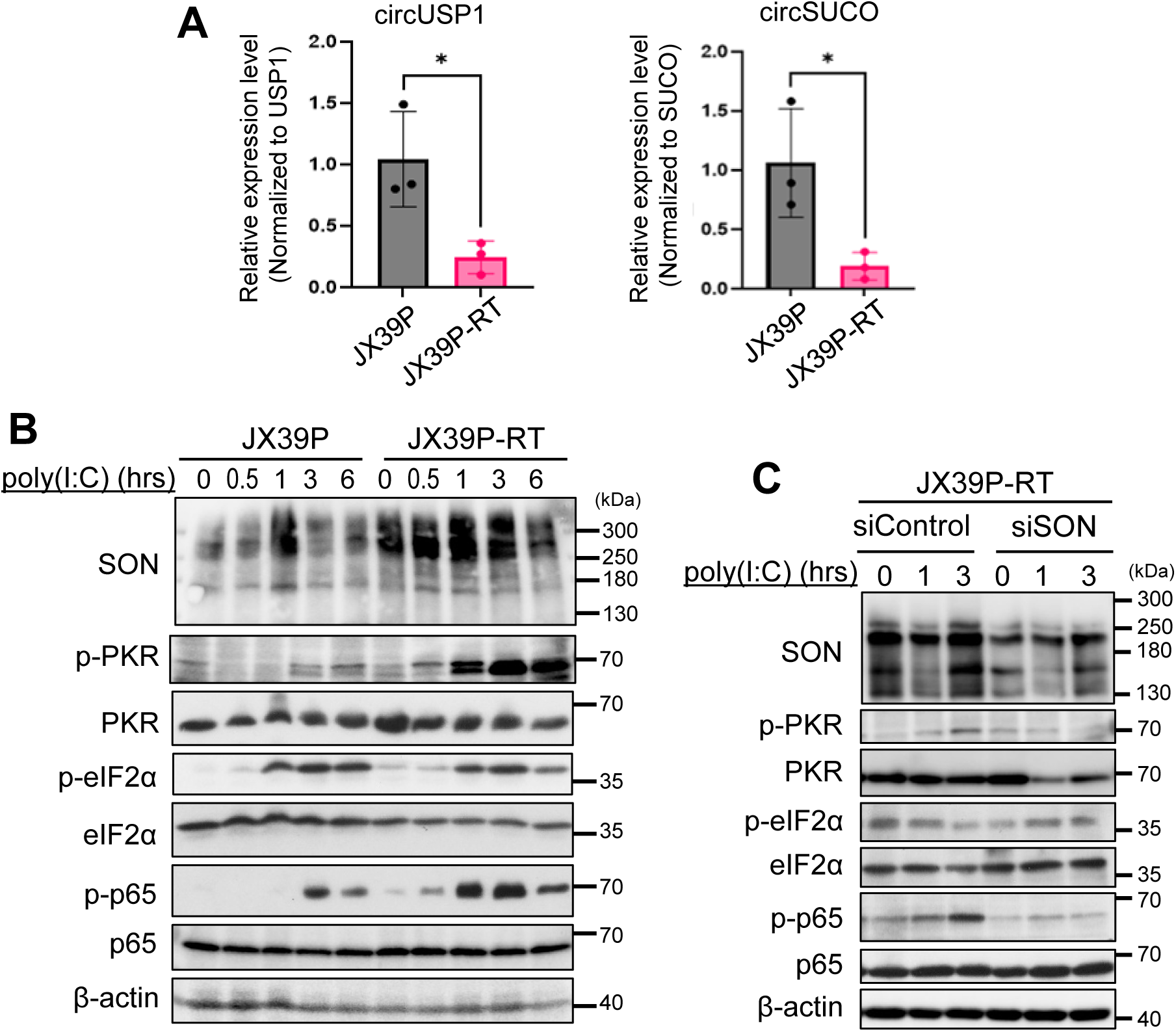
Elevated SON contributes to enhanced PKR/NFκB signaling in radiation-resistant GBM. **(A)** qRT-PCR using divergent primers showing reduced expression of circUSP1 and circSUCO in JX39P-RT compared with JX39P cells. CircRNA expression was normalized to the corresponding linear parental transcript. \**p* < 0.05. **(B)** Western blot analysis of JX39P and JX39P-RT cells following poly(I:C) stimulation for the indicated times, showing enhanced PKR phosphorylation and downstream NFκB activation in JX39P-RT cells. **(C)** Western blot analysis of JX39P-RT cells transfected with control siRNA (siCtrl) or SON-targeting siRNA (siSON) followed by poly(I:C) stimulation. SON depletion attenuated phosphorylation of PKR and p65 in JX39P-RT cells.

We next examined whether the acquired radiation-resistant state is accompanied by altered PKR signaling. Following poly(I:C) stimulation, JX39P-RT cells exhibited earlier and more sustained PKR phosphorylation compared with parental JX39P cells (**Figure 5B**). Enhanced PKR activation in JX39P-RT was accompanied by increased phosphorylation of p65, indicating increased NFκB activation. Together, these results demonstrate enhanced activation of the PKR/NFκB signaling pathway in the radiation-resistant JX39P-RT cells.

To determine whether elevated SON contributes to this enhanced PKR/NFκB signaling, we depleted SON in JX39P-RT cells and examined the response to poly(I:C). SON knockdown markedly reduced PKR phosphorylation and downstream p65 phosphorylation following poly(I:C) stimulation (**Figure 5C**). Thus, elevated SON expression contributes to the enhanced PKR/NFκB response in JX39P-RT cells.

Together, these findings show that radiation-resistant JX39P-RT cells exhibit suppression of specific SON-regulated circRNAs together with enhanced PKR/NFκB activation. Depletion of SON attenuates this enhanced signaling, positioning SON upstream of PKR/NFκB activation in radiation-resistant GBM and supporting a model in which elevated SON contributes to the stress-response phenotype associated with radiation resistance.

### Overexpression of SON-regulated circRNAs suppresses PKR activation

To directly test whether SON-regulated circRNAs affect PKR signaling, we generated circRNA overexpression constructs by inserting the circRNA-forming exons, together with 150 bp of upstream and downstream intronic sequence, into a vector containing inverted repeats to promote circularization (**Figure 6A**).

**Figure 6.**
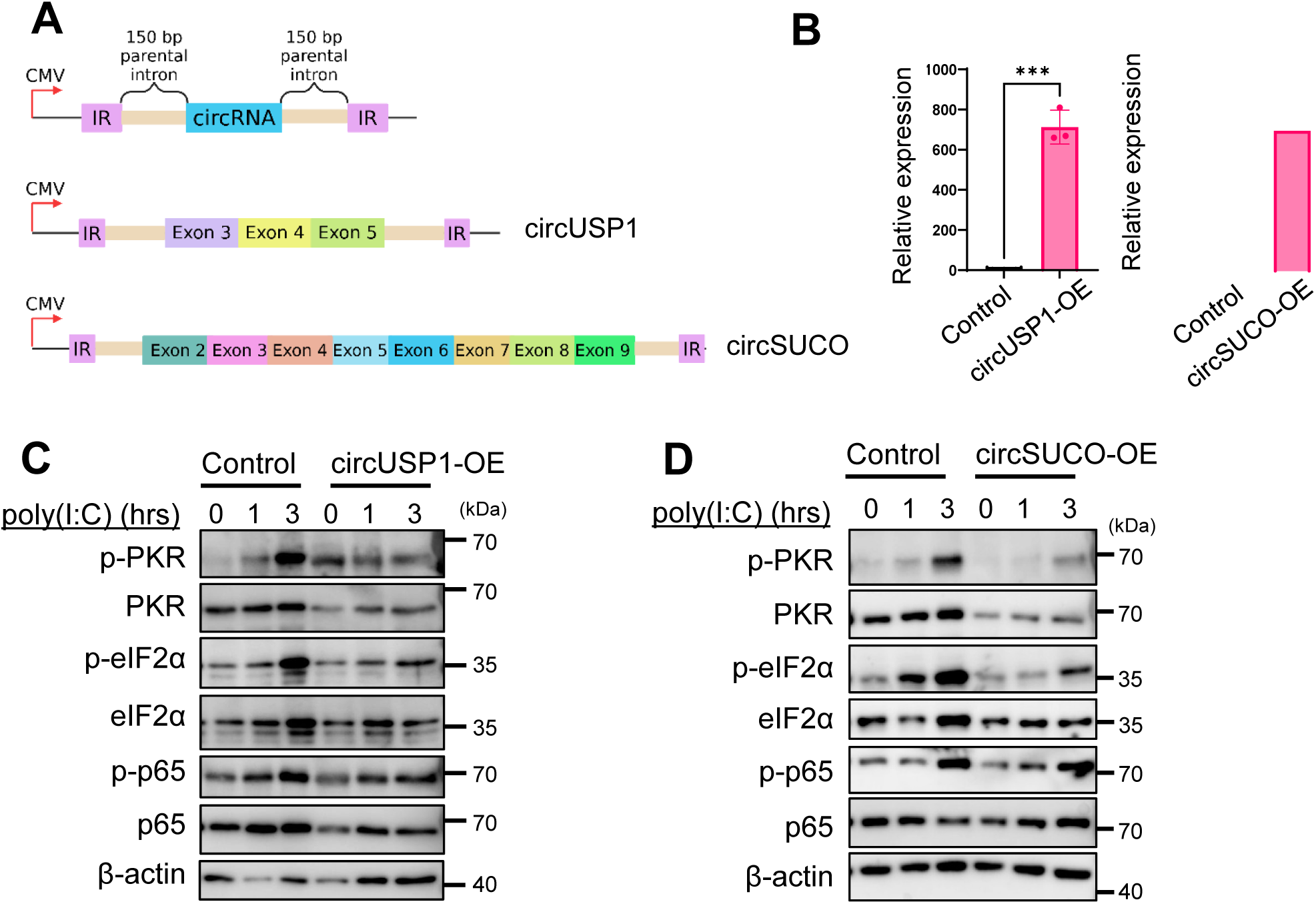
Overexpression of SON-suppressed circRNAs attenuates PKR activation in radiation-resistant GBM. **(A)** Schematic of circUSP1 and circSUCO overexpression constructs containing the circRNA-forming exons flanked by 150 bp of parental intronic sequence and inverted repeats (IR) to promote circularization. **(B)** qRT-PCR validation of circUSP1 and circSUCO overexpression in LN229RR cells. circUSP1 expression was measured in untreated RNA and normalized to YWHAZ because of its low endogenous abundance, whereas circSUCO expression was measured following RNase R treatment and normalized to circHIPK3. \*\**p* < 0.01; \*\*\**p* < 0.001. **(C)** LN229RR cells were transfected with circUSP1 overexpression vector (circUSP1-OE) or empty vector (Control) and stimulated with poly(I:C) for 0, 1, or 3 h. circUSP1 overexpression attenuated PKR phosphorylation and downstream p65 and eIF2α phosphorylation following poly(I:C) stimulation. **(D)** LN229RR cells were transfected with circSUCO overexpression vector (circSUCO-OE) or empty vector (Control) and stimulated with poly(I:C) for 0, 1, or 3 h. circSUCO overexpression attenuated PKR and eIF2α phosphorylation, with little effect on p65 phosphorylation.

We confirmed overexpression of circUSP1 and circSUCO by qRT-PCR (**Figure 6B**). Because endogenous circUSP1 expression is extremely low in radiation-resistant GBM cells and was difficult to detect following RNase R treatment, circUSP1 expression was measured in untreated RNA and normalized to *YWHAZ*. For circSUCO, expression was measured in RNase R-treated samples and normalized to circHIPK3, which was unchanged in our circRNA-sequencing datasets. Both circRNAs were significantly increased following transfection of their respective overexpression constructs.

We next examined whether restoration of these circRNAs affects PKR activation following poly(I:C) stimulation. Overexpression of either circUSP1 or circSUCO reduced PKR phosphorylation (**Figure 6C** and **D**), recapitulating the reduced PKR activation observed following SON depletion. circUSP1 also reduced p65 phosphorylation, whereas circSUCO had little effect on NFκB activation despite reducing PKR phosphorylation. Both circRNAs reduced eIF2α phosphorylation.

These results show that restoring circUSP1 or circSUCO is sufficient to suppress PKR activation in GBM cells. The differential effects on downstream NFκB signaling also suggest that individual circRNAs may not regulate all components of the PKR stress-response pathway in the same manner.

### Overexpression of SON-suppressed circRNAs reverses radiation resistance in GBM

We first examined whether restoration of SON-suppressed circRNAs affects GBM cell proliferation. Overexpression of either circUSP1 or circSUCO significantly increased the percentage of Ki-67-positive LN229 cells compared with control cells (**Figure 7A and B**). Given the relationship between proliferative state and radiation response, this finding prompted us to determine whether restoring these circRNAs could alter the response of GBM cells to irradiation.

**Figure 7.**
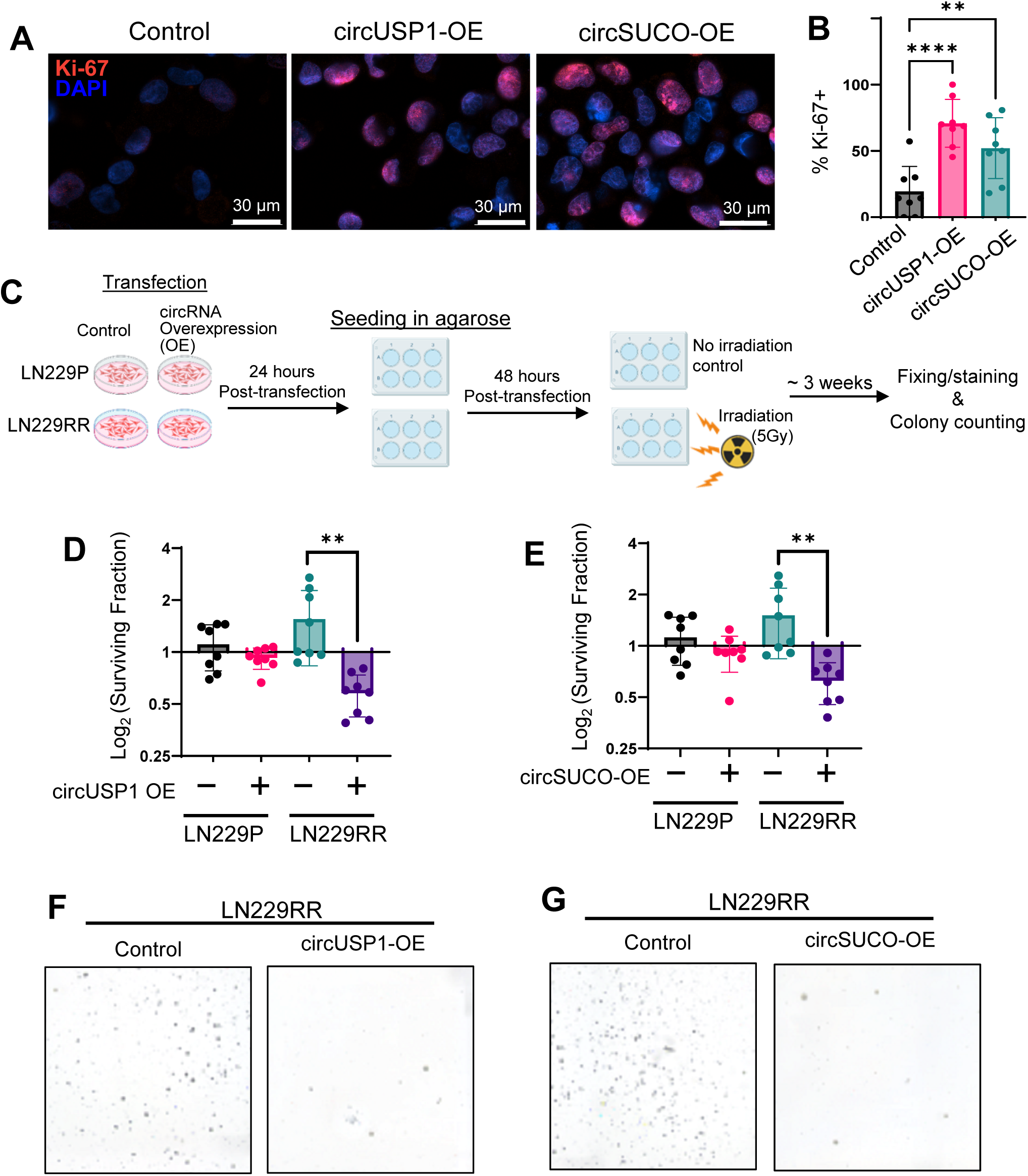
Overexpression of circUSP1 and circSUCO increases the proliferative fraction and reduces clonogenic survival in radiation-resistant GBM cells. **(A)** Representative immunofluorescence images of Ki-67 staining in LN229 cells transfected with empty vector (Control), circUSP1-OE, or circSUCO-OE. Ki-67 is shown in red and nuclei are counterstained with DAPI (blue). **(B)** Quantification of the percentage of Ki-67-positive cells. Overexpression of circUSP1 or circSUCO significantly increased the percentage of Ki-67-positive cells compared with control cells. Eight fields of view were analyzed per condition. **(C)** Schematic of the 3D clonogenic assay. LN229P and LN229RR cells were transfected with circUSP1-OE, circSUCO-OE, or empty vector, embedded in agarose 24 h after transfection, and irradiated (5Gy) 48 h after transfection. Colonies were stained and counted at the experimental endpoint. **(D)** circUSP1 overexpression significantly reduced the surviving fraction of LN229RR cells compared with empty vector controls, whereas no significant difference was observed in LN229P cells. **(E)** circSUCO overexpression significantly reduced the surviving fraction of LN229RR cells compared with empty vector controls, whereas no significant difference was observed in LN229P cells. **(F)** Representative images of colonies formed by LN229RR cells transfected with empty vector or circUSP1-OE. **(G)** Representative images of colonies formed by LN229RR cells transfected with empty vector or circSUCO-OE. \*\**p* < 0.01; \*\*\*\**p* < 0.0001

To directly test this, we performed a 3D clonogenic assay using parental LN229P cells and the matched radiation-resistant LN229RR cells (**Figure 7C**). Cells were transfected with circUSP1, circSUCO, or a control vector, embedded in agarose, and exposed to irradiation. Importantly, overexpression of either circUSP1 or circSUCO significantly reduced the surviving fraction of LN229RR cells compared with control cells (**Figure 7D - G**). In contrast, neither circRNA significantly altered the surviving fraction of parental LN229P cells. Thus, the effect of circUSP1 and circSUCO restoration was most evident in cells that had acquired radiation resistance.

Together, these findings show that overexpression of SON-suppressed circUSP1 and circSUCO selectively resensitizes GBM cells that have acquired radiation resistance. These results identify SON-regulated circRNAs as functional mediators of radiation response and suggest that increasing the abundance of PKR-inhibitory circRNAs may represent a potential strategy for radiosensitizing resistant GBM.

## Discussion

In this study, we identify SON as a previously unrecognized regulator of circRNA biogenesis in GBM and define a SON/circRNA/PKR signaling axis associated with radiation resistance. Although SON is known to regulate alternative splicing and oncogenic RNA processing in GBM, its role in backsplicing and circRNA formation has not been established. We show that SON depletion produces widespread increases in circRNA abundance and attenuates PKR signaling, whereas radiation-resistant GBM cells exhibit the opposite pattern, with increased SON expression, widespread circRNA loss, and enhanced PKR activation. Importantly, overexpression of the SON-suppressed circRNAs circUSP1 and circSUCO inhibits PKR activation and selectively resensitizes radiation-resistant GBM cells to irradiation. These findings identify circRNA loss as a functional component of the radiation-resistant phenotype and connect SON-dependent RNA processing with stress signaling and radiation response.

CircRNAs are highly enriched in the nervous system and generally accumulate during neuronal differentiation and maturation. Consistent with previous studies, circRNA abundance is markedly reduced in GBM compared with normal brain, but the mechanisms underlying this loss remain poorly understood. Our findings identify SON as a regulator of circRNA abundance in GBM. Depletion of SON resulted in widespread increases in circRNAs detected independently by two circRNA-analysis pipelines, indicating that SON normally restricts circRNA formation. Given the established role of SON in spliceosome recruitment and efficient processing of transcripts containing weak splice sites, one possibility is that elevated SON favors canonical splice-site utilization, thereby reducing opportunities for backsplicing. The molecular basis of this effect remains to be determined, but our findings extend the function of SON beyond regulation of alternative linear splicing and suggest that SON also influences the balance between linear and circular RNA production.

Previous studies have shown that endogenous circRNAs contain short, imperfect dsRNA structures that can bind to and inhibit PKR. Our findings extend this mechanism to GBM and identify SON as an upstream regulator of this pathway. SON knockdown increased circRNA abundance and attenuated PKR activation, suggesting a relationship between SON-dependent circRNA suppression and PKR signaling. Importantly, overexpression of individual SON-suppressed circRNAs provided functional evidence for this connection. Overexpression of either circUSP1 or circSUCO reduced PKR phosphorylation following poly(I:C) stimulation, recapitulating the effect observed following SON depletion. These findings demonstrate that specific circRNAs suppressed by SON are capable of inhibiting PKR activation in GBM cells. Thus, elevated SON expression may enhance PKR signaling by reducing the abundance of circRNAs that normally restrain PKR activation. This finding gives functional significance to circRNA loss in GBM and suggests that circRNAs can directly influence stress-response signaling rather than simply reflecting the differentiation state of tumor cells.

The relationship between SON, circRNA abundance, and PKR signaling became more pronounced following the acquisition of radiation resistance. Radiation-resistant cells exhibited increased SON expression, and JX39P-RT cells showed widespread circRNA loss together with enhanced PKR/NFκB activation. circUSP1 and circSUCO were also reduced in JX39P-RT cells, while SON depletion attenuated PKR and NFκB activation in these cells. Most importantly, overexpression of circUSP1 or circSUCO suppressed PKR activation and significantly reduced clonogenic survival following irradiation in radiation-resistant LN229RR cells, while having little effect on the radiation response of parental LN229P cells. This selective radiosensitization indicates that increasing the abundance of these circRNAs can counter, at least in part, the acquired radiation-resistant phenotype. Interestingly, circUSP1 and circSUCO overexpression also increased the percentage of Ki-67-positive GBM cells. Because proliferative state can influence radiation response, this change may contribute to radiosensitization; however, whether increased proliferation is required for this effect remains to be determined. Together, these findings connect SON-dependent circRNA loss and enhanced PKR signaling with radiation resistance and suggest that increasing the abundance of specific PKR-inhibitory circRNAs may provide a strategy for resensitizing resistant GBM to irradiation.

Despite the significance of these findings, several limitations should be considered. First, the functional studies were performed primarily in cell-based models, and *in vivo* studies will be necessary to determine whether overexpression of SON-regulated circRNAs can enhance radiation response in orthotopic GBM models. Such studies will also be important for determining whether therapeutic delivery of circUSP1 or circSUCO can achieve sufficient and sustained expression within GBM tumors to enhance the response to radiotherapy. Although our findings identify elevated SON as an upstream contributor to circRNA suppression and PKR activation, directly targeting SON may have limited therapeutic feasibility because of its essential roles in RNA processing and development. Indeed, SON haploinsufficiency causes ZTTK syndrome, a multisystem neurodevelopmental disorder, underscoring the potential consequences of reducing SON activity in normal tissues(40). Targeting downstream components of this pathway, particularly restoration of specific PKR-inhibitory circRNAs, may therefore provide a more selective approach to modulating this pathway in radiation-resistant tumors. Second, although our data identify SON as a regulator of circRNA biogenesis, the molecular mechanism by which SON suppresses backsplicing remains to be determined. Future studies examining SON occupancy at circRNA-producing transcripts and its interactions with spliceosomal components may clarify how SON regulates the balance between canonical splicing and circRNA formation. Finally, circUSP1 and circSUCO represent only two members of a much broader population of SON-regulated circRNAs. Additional circRNAs may contribute to PKR regulation and radiation response, and defining the circRNAs with the strongest functional and therapeutic potential will be important for future studies.

In conclusion, our study identifies a previously unrecognized SON/circRNA/PKR regulatory pathway that links RNA processing to radiation resistance in GBM. We show that SON suppresses circRNA biogenesis, that SON-regulated circRNAs restrain PKR signaling, and that this regulatory relationship is altered in radiation-resistant GBM, where increased SON expression is accompanied by widespread circRNA loss and enhanced PKR activation. Most importantly, overexpression of the SON-suppressed circRNAs circUSP1 and circSUCO suppresses PKR activation and selectively resensitizes radiation-resistant GBM cells to irradiation. These findings identify specific circRNAs not only as functional regulators of radiation response but also as potential therapeutic tools for restoring radiation sensitivity in resistant GBM. By connecting SON-dependent RNA processing, circRNA-mediated regulation of stress signaling, and acquired radiation resistance, our work reveals a previously unrecognized mechanism of therapeutic resistance and identifies increasing the abundance of specific PKR-inhibitory circRNAs as a potential strategy for resensitizing radiation-resistant GBM to radiotherapy.

## Materials and Methods

### Cell culture and GBM models

Normal human astrocytes (NHA), LN229, MGG18, and HEK293T cells were maintained in DMEM High Glucose (Elabscience, PM150210) supplemented with 10% FBS (GeminiBio, S11150). Cells were dissociated using 0.25% Trypsin-EDTA (Sigma Aldrich, T4049) and cryopreserved in 90% FBS and 10% DMSO. Glioma stem cells (GSC83)(29) and patient-derived xenolines (JX14P and JX39P/P-RT) were cultured in DMEM/F12 (Elabscience, PM150310) supplemented with Gem21 NeuroPlex (GeminiBio, 400-161-010), EGF (20 ng/mL; Sino Biological, 10605-HNAE), bFGF (20 ng/mL; Sino Biological, 10014-HNAE), and heparin (5 µg/mL; MedChemExpress, HY-17567C). Cells were dissociated using Accutase (Innovative Cell Technologies, AT104) and cryopreserved in CellBanker2 (Amsbio, 11914). All cultures were maintained at 37°C and 5% CO_2_.

### Generation of radiation-resistant models

Radiation-resistant LN229 cells (LN229RR) were generated by exposing parental LN229 cells to 5 Gy every 4 days for four cycles. To maintain resistance, LN229RR cells were re-irradiated with 5 Gy every fourth passage. The radiation-resistant GBM PDX line JX39P-RT was generated, as described previously (39), by subcutaneous implantation of freshly resected GBM tissue into athymic nude mice followed by repeated cycles of fractionated irradiation (2 Gy, three times per week for two weeks; total 12 Gy).

### shRNA-mediated SON knockdown

Stable SON knockdown in GSC83 cells was achieved using pLL3.7 lentiviral vectors containing shSON constructs targeting exon 3. Lentivirus was generated in HEK293T cells using third-generation packaging plasmids (pMDLg/pRRE, pRSV-Rev, pMD2.G) and PEI (1:3 DNA:PEI). Viral supernatant was collected at 48 h, filtered (0.45 µm), and concentrated by ultracentrifugation. GSC83 cells were infected by adding viral supernatant directly to media, followed by puromycin selection (1 µg/mL) beginning 24 h post-infection. RNA was harvested 72 hours post-infection.

### siRNA-mediated SON knockdown

LN229, JX39P, and JX39P-RT cells were transfected with SON-targeting siRNA (Thermo Fisher, Custom Silencer Select) or non-targeting control siRNA (Thermo Fisher, 4390844). For LN229, cells were seeded at 200,000 cells/well and transfected using RNAiMAX (Invitrogen, 13778075) according to manufacturer instructions. For JX39P and JX39P-RT, reverse transfection was performed by pre-forming RNAiMAX/siRNA complexes in wells prior to cell addition. Cells were harvested 48 h post-transfection.

### CircRNA overexpression

Gene fragments containing the circRNA-forming exons of circSEC24A, circUSP1, and circSUCO together with 150 bp of upstream and downstream flanking intronic sequence were synthesized by Twist Bioscience with restriction sites for SacII and ClaI. Inserts and vector (Addgene 60648) were digested with SacII (New England Biolabs, R0157) and ClaI (New England Biolabs, R0197) and ligated using T4 DNA ligase (New England Biolabs, M0202). LN229 cells were transfected using PEI (1:3 DNA:PEI) and harvested at 48 hours. Overexpression was confirmed using divergent primers.

### RNA extraction, cDNA synthesis, and qRT-PCR

RNA was isolated using TRIzol (Invitrogen, 15596018) followed by DNaseI treatment (Invitrogen, AM1906). RNA purity was assessed by NanoDrop (A260/280 > 1.7). RNase R digestion was performed using 3–5 µg RNA with RNase R (NEB, M0100S) at 1 U/µg RNA for 30 minutes at 37°C, followed by purification using miRNeasy (Qiagen, 217004). cDNA synthesis was performed using SuperScript III (Invitrogen, 18-080-051). qRT-PCR was performed using iTaq SYBR Green Supermix (Bio-Rad, 1725124) on a Bio-Rad CFX Connect. Linear transcripts were quantified using convergent primers; circRNAs were quantified using divergent primers. Melt-curve analysis confirmed specificity. Expression was normalized to YWHAZ, the corresponding linear parental transcript, or circHIPK3, as indicated for each experiment.

### Sequencing analysis

GSC83 RNase R-treated libraries were analyzed using CIRIquant v1.1.3 with GRCh38 and Ensembl release 111. CircRNAs were detected de novo using the integrated CIRI2 module, and backsplice junction (BSJ) and forward-splice junction (FSJ) reads were quantified (30, 32). Matched untreated libraries were processed using CIRIquant’s RNase R-correction mode to estimate circRNA abundance. Count matrices and circRNA annotations were generated using prep_CIRIquant, and gene-level counts from StringTie were used for normalization. Differential expression was performed using CIRI_DE_replicate (TMM normalization; edgeR GLM).

As an orthogonal approach, RNase R-treated reads were aligned using STAR with chimeric-read detection enabled (41). Candidate BSJs were identified using CIRCexplorer2, filtered based on junction support (31), and re-quantified using CIRIquant. Differential expression was assessed using CIRI_DE_replicate.

CIRIquant outputs were analyzed in R. CircRNAs were assigned to host genes using CIRIquant GTF annotations. CircRNAs with log_2_fold change > |1| and p < 0.05 were considered altered. Volcano plots were generated from CIRI_DE_replicate results, and heatmaps were constructed from circular-to-linear junction ratios.

CircRNAs from JX39P and JX39P-RT libraries were identified and quantified using CIRIquant with CIRI2 for de novo BSJ detection. BSJ/FSJ counts, junction ratios, and circRNA abundance estimates were generated using prep_CIRIquant, and differential expression was performed as described above.

Gene set enrichment analysis (GSEA) was performed on circRNA host genes using a pre-ranked approach in clusterProfiler with gene sets obtained from msigdbr (42). If multiple circRNAs mapped to the same gene, the circRNA with the largest absolute log_2_fold change was used. Enrichment was tested against Hallmark and GO Biological Process collections (gene-set size 10-500). Benjamini–Hochberg correction was applied, and gene sets with adjusted p < 0.25 were considered exploratory.

Enrichment analysis was conducted against the MSigDB Hallmark, Gene Ontology Biological Process, and Reactome collections using gene-set sizes between 10 and 500 genes. Multiple-testing correction was performed using the Benjamini-Hochberg method. Gene sets with adjusted p < 0.25 were considered exploratory enrichment signals. Positive normalized enrichment scores indicate enrichment among host genes associated with circRNAs that increased following SON knockdown, whereas negative scores indicate enrichment among host genes associated with circRNAs that decreased following SON knockdown.

### Western blotting

Cells were lysed in 2× SDS sample buffer and sonicated. Lysates were clarified by centrifugation at 13,000 × *g* for 30 min at 4°C. Samples were resolved by SDS-PAGE, transferred to PVDF membranes, and probed with antibodies: SON (custom rabbit polyclonal antibody raised against an epitope corresponding to amino acids 74-88 of human SON), PKR (Invitrogen, 700286), p-PKR (Abcam, AB81303), eIF2α (Protein Tech, 11170), p-eIF2α (Cell Signaling, 3398), p65 (Santa Cruz Biotechnology, sc-8008), p-p65 (Cell Signaling, 3033), and β-actin (Protein Tech, 66009). Blots were imaged using chemiluminescence.

### Poly(I:C) stimulation

Poly(I:C) (MedChemExpress, HY-135748) was delivered using RNAiMAX. LN229 cells received 10 µg/mL poly(I:C) via forward transfection; LN229 and JX39P/JX39P-RT cells were stimulated via reverse transfection. Cells were harvested at indicated time points.

### Irradiation

Cells were exposed to 5 Gy, unless otherwise indicated, using an X-ray irradiator at a dose rate of 1.5 Gy/min. Irradiated cells were used for Ki-67 immunofluorescence or clonogenic survival assays.

### Immunofluorescence and proliferation analysis

Cells were fixed and permeabilized in ice-cold methanol (−20°C, ≥20 min), blocked in PBS containing 2% BSA and 0.1% Triton X-100, and stained with antibodies against SON (custom rabbit polyclonal antibody raised against an epitope corresponding to amino acids 74 - 88 of human SON) or Ki-67 (DSHB, AFFN-KI67-3E6), as indicated. Alexa Fluor-conjugated secondary antibodies were applied, and nuclei were counterstained with DAPI. Images were acquired using a Lionheart FX automated imager. For Ki-67 analysis, the proliferative fraction was determined as the percentage of Ki-67-positive cells among total DAPI-positive cells.

### Clonogenic assay

Clonogenic assays were performed according to a published protocol (43). LN229P and LN229RR cells were transfected with circRNA expression vectors. At 24 hours post-transfection, cells were embedded in a two-layer agarose system consisting of a 0.75% agarose base layer and a 0.375% agarose top layer containing 1,000 - 5,000 cells. At 48 hours post-transfection, cultures were irradiated with 5 Gy or left unirradiated (0 Gy) and maintained with media changes every 3 - 4 days. At the experimental endpoint, colonies were fixed with PFA, stained with toluidine blue, and counted. Plating efficiency (PE) was calculated as: 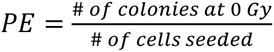. Surviving fraction (SF) is calculated as: 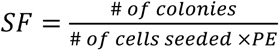

### CircRNA secondary structure prediction

Secondary structure prediction was performed using RNAfold (ViennaRNA Package) with circular topology enabled. Minimum free energy structures and base-pair probabilities were analyzed to identify internal dsRNA regions according to criteria described by Liu et al.

### Statistics

Statistical analyses were performed using GraphPad Prism and R. qRT-PCR data were analyzed using the ΔΔCt method, with technical triplicates averaged to generate a single value per biological replicate. Unless otherwise indicated, comparisons between two groups were made using unpaired two-tailed Student’s t-tests. Multi-group experiments, including irradiation dose– response and clonogenic assays, were analyzed using one-way or two-way ANOVA with appropriate post hoc correction. CircRNA-sequencing differential expression was assessed using edgeR-based generalized linear modeling with TMM normalization and Benjamini–Hochberg correction. Statistical significance was defined as p < 0.05.

### Figure Preparation

Figures were generated using GraphPad Prism, R, or BioRender.

## Acknowledgments

This work was supported by the National Institutes of Health (NIH) (R01CA236911 and R01HL168659 to E.E.A.; R01NS138515 to S.O.; R01HL158800 and R01HL158875 to S.S.L.), the Brain Tumour Charity (to S.O.), the University of Alabama at Birmingham (UAB) Heersink School of Medicine, UAB Department of Pathology, and UAB O’Neal Cancer Center’s O’Neal Invests Program (Catalyst grant to E.E.A. and C.D.W.). S.D. and C.A.H. were supported by the NIH Training Program T32 (grant no. T32GM135028 for S.D.; grant no. T32CA047888 for C.A.H.). This content is solely the responsibility of the authors and does not necessarily represent the views of the NIH.

